# Towards region-specific propagation of protein functions

**DOI:** 10.1101/275487

**Authors:** Da Chen Emily Koo, Richard Bonneau

## Abstract

**Motivation:** Due to the nature of experimental annotation, most protein function prediction methods operate at the protein-level, where functions are assigned to full-length proteins based on overall similarities. However, most proteins function by interacting with other proteins or molecules, and many functional associations should be limited to specific regions rather than the entire protein length. Most domain-centric function prediction methods depend on accurate domain family assignments to infer relationships between domains and functions, with regions that are unassigned to a known domain-family left out of functional evaluation. Given the abundance of residue-level annotations currently available, we present a function prediction methodology that automatically infers function labels of specific protein regions using protein-level annotations and multiple types of region-specific features.

**Results:** We apply this method to local features obtained from InterPro, UniProtKB and amino acid sequences and show that this method improves both the accuracy and region-specificity of protein function transfer and prediction by testing on both human and yeast proteomes. We compare region-level predictive performance of our method against that of a whole-protein baseline method using a held-out dataset of proteins with structurally-verified binding sites and also compare protein-level temporal holdout predictive performances to expand the variety and specificity of GO terms we could evaluate. Our results can also serve as a starting point to categorize GO terms into site-specific and whole-protein terms and select prediction methods for different classes of GO terms.

**Availability:** The code is freely available at: https://github.com/ek1203/region_spec_func_pred

## 1. Introduction

Proteins are involved in nearly every cellular process and function, including cell organization, biochemical catalysis, signaling and transport. The exponentially large number of possible sequence combinations enables proteins to exhibit the necessary diverse sequential, structural and functional properties required by the cell. Protein function is often defined or modified by specific interactions with other molecules, therefore knowing what, where and how proteins interact is important in elucidating the cellular machinery of life [1]. Proteins can be multifunctional by having completely different functions in different contexts (“moonlighting proteins” like crystallins [44, 24]), by binding to multiple substrates and catalyzing multiple reactions (“promiscuous proteins” [23, 30]) or by having combinations of domains in different sequential orders [5]. Comprehensive experimental characterization of protein function is thus a laborious, expensive and time-consuming process, which is especially true for proteins found in non-model and multicellular organisms.

With the advent of the Gene Ontology (GO) project [4], computational prediction of protein function has become more viable and it is an ongoing quest to improve the number and quality of predictions. Many different approaches have been proposed over decades [26, 57, 33, 37, 54, 40, 56, 49, 12, 13] and reviewed extensively [9, 48, 34, 8, 31].

Although not an all-encompassing evaluation, results from large-scale Critical Assessment of protein Function Annotation (CAFA) experiments [45, 27] and related blind tests like MouseFunc [42] offer important insights into the performances of state-ofthe-art computational protein function prediction methods.

The underlying principle of protein function prediction is the transfer of function from a known protein to a query protein based on shared features. Such features include protein sequence, structure and pairwise associations from high-throughput experimental data like protein-protein interaction and gene co-expression. BLAST [3], for example, is one of the most widely used sequence-based tools and is regularly used as a benchmark to evaluate the performances of more complex methods. Due to the nature of experimental annotation (and the prevalence of genetics as a means of connecting genes to functions), most protein function prediction methods operate at the whole-protein level (i.e. functions are transferred directly between whole-chain proteins). However, the majority of proteins are composed of one or more structural and functional units called domains [20], which can function independently or in combination with other domains (“supra-domains” [59]).

Ideally, functions wholly encapsulated within such regions should be confined and uncoupled from other regions of the protein in a region-specific annotation scheme. This is especially true for GO terms in the Molecular Function (MF) branch of the ontology. Currently, a few such annotated domain-centric resources exist through a combination of manual and automated curation (e.g. Pfam [18], SUPERFAMILY [22], CATH-GENE3D [55], meta-database InterPro [17]), allowing functions to be transferred to novel proteins assigned with known and annotated domain families. Other domain-centric methods have also been described previously [53, 19, 36, 16, 47, 14, 13] to automatically associate functions directly to domain families before integrating them for protein function prediction.

These resources are generally very sparse as they require a fine balance between sufficient coverage of the domain space and the applicability of the annotations to all proteins matching the given domain signature [10]. This is especially problematic for large and diverse families, and hinders the mapping of specific GO terms. Another common weakness of these domain-centric approaches is that they depend entirely on predicted domain family assignments, which not only differ based on different classification and identification schemes, but are also constantly changing and updating [51]. Additionally, these predicted assignments only cover about a third of the total residues in the proteomes. For example, even though approximately 55% of yeast and 68% of human protein sequences (UniProtKB [6] reference proteomes release 2017_10) have at least one ’DOMAIN’ entry type assigned (InterPro database [17] release 65.0, Oct 2017), less than 32% and 38% of total residues in each proteome, respectively, are actually covered by the assignments. This can be due to the fact that the majority of domain families identified are structured domains. Intrinsically disordered regions, which are prevalent in eukaryotic genomes and have been established to actively participate in diverse protein functions [58], are excluded entirely from functional evaluation. In addition, the treatment of domains as binary features of proteins prevent the transfer of function from annotated to unannotated domain families, which can have shared functions as well.

Therefore, we find it pertinent to decompose proteins into regions *containing* and *not containing* domain family assignments, and then build a method that can transfer protein labels at the region level explicitly (including regions not covered by traditional domain assignments). For example, instead of representing “BAI1-associated protein 2like protein 1” (*BAIAP2L1*, UniProtKB accession: Q9UHR4) as a 511 residue protein with two domains, an IMD/I-BAR domain (residues 1 to 249) and an SH3 domain (residues 339 to 402), we will represent this protein as four regions, with regions 1 and 3 containing the well-annotated domains domains and the remaining regions 2 (residues 250 to 338) and 4 (residues 403 to 511) containing (prior to prediction) no assigned domain families. These unassigned regions will still contain sequence information (and sequence derived features) and other site-specific feature annotations, such as post-translational modifications and F-actin binding sites, from databases with manual curation like UniProtKB [6] that can provide functional clues (features).

Here, we detail our approach to generating protein regions using curated site-specific features and to localizing known protein function labels to these regions automatically based on related approaches to structured sentiment analysis [32]. We evaluate the prediction accuracy of our region-specific framework for a variety of GO terms at both regionand protein-levels for yeast and human proteomes and describe performance improvements over using a whole-protein baseline model. Our results show that many GO terms benefit from applying this region-specific framework and that different types of GO terms should be treated differently in function prediction pipelines depending on their extent of functional localization.

## 2. Methods

The general process outline of our method is to: (1) split protein sequences into potential functional regions based on the presence and absence of domain and protein family assignments, (2) encode the regions as separate feature vectors based on data sources summarized in Table 1, and (3) train model to infer region function labels from known protein labels. This is summarized in Figure 1.

**Figure 1:**
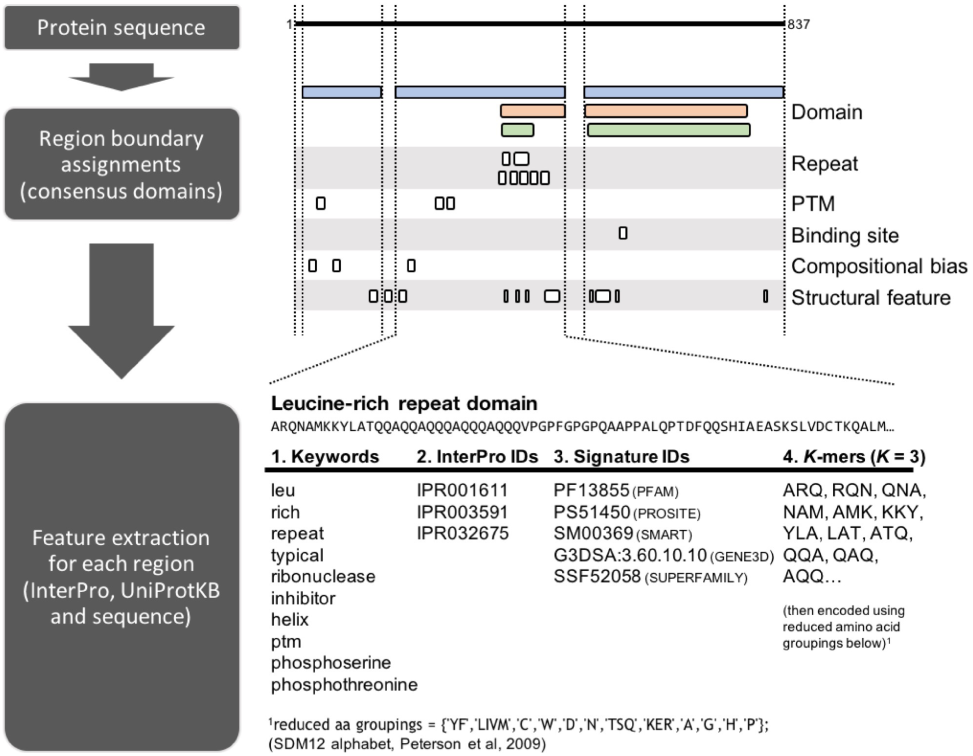
Diagram of processing pipeline using protein Glucose-repressible alcohol dehydrogenase transcriptional effector (P31384) as an example. Features assigned to the protein sequence are grouped by feature types labeled on the right and assignments from different databases for the same type are shown on separate rows. Region boundaries are delineated by vertical lines based on consensus domain assignments from different databases (3 in this case) and different feature annotations from InterPro and UniProtKB within each region are parsed into the 4 major feature types shown in the table at the bottom.

**Table 1:** Data sources

| Data type | Version |
| --- | --- |
| Protein set | UniProtKB Reference Proteomes release 2017_10 |
| GO annotations (protein) | UniProt-GOA release 145 (2015-06) and 171 (2017-10) [non-IEA only <sup>1</sup>] |
| GO annotations (domain/region) | external2go (2015-07-25 and 2017-11-14), inferred from binding sites (NBench and BioLiP) |
| InterPro features | InterProScan 5.26-65.0 |
| UniProt features | UniProtKB/Swiss-Prot Release 2017_10 |
| DisProt features | DisProt 7 release 0.5 |

<sup>1</sup>Annotations from all evidence codes except 'Inferred from Electronic Annotation'.

### 2.1. Generating region boundaries

The first step in this approach is to split the protein sequences into potential functional regions. To do that, we processed the amino acid sequences with InterProScan [28] and used the feature annotations to build consensus region boundaries. InterPro entry types used include Domain, Family, Homologous superfamily and select unintegrated signatures from InterPro [17], Signal peptide, Transmembrane and Non-transmembrane from PHOBIUS [29], and Disorder from MobiDB-lite [41]. All Repeats and Sites annotations are excluded during this process. Aside from Signal peptide, which is included whenever available, initial region boundaries are assigned exclusively by types in the following order of precedence: Domain, Family and Homologous superfamily, Unintegrated signatures, Transmembrane, Non-transmembrane and Disorder.

Unassigned **terminal regions** less than 18 residues in length (selected based on signal length distributions [7]) are merged with immediate neighboring regions to remove excessive numbers of short peptides without losing potential targeting and retention signal peptides, while longer terminal regions are retained as separate regions. **Interdomain regions** less than 20 residues in length (selected based on linker length distribution [11]) are also discarded to remove the majority of linker regions.

### 2.2. Data representation and feature generation

Each protein/region is represented as a fixedlength vector in four different feature spaces encoded with the data types listed below.

1. ***K*-mers** a collection of consecutive, overlapping, *k* -residue long sub-sequences of the sequence of the region itself. For example, a sequence of ’ABCDE’ will result in 3-mers of ’ABC’, ’BCD’ and ’CDE’. The length of 3 was tested to get a compromise between capturing sufficient protein fold information and having a feature vector that is not too large to deal with. In addition, the primary sequence was encoded using a reduced amino acid alphabet (SDM12 as described and compared in [43]), where similar amino acids are clustered together to give 12 groups instead of the initial 20. This was done to increase sensitivity to regions with structural similarity, which can be more important for some functions than sequence identity.
2. **Keywords** a vocabulary of individual words parsed from descriptions of features assigned from UniProtKB, InterPro entries, original member databases and DisProt. For example, a region assigned with feature ’WD repeat-containing’ will contain keywords ’WD’, ’repeat’ and ’containing’, allowing the region to have a non-zero similarity score when compared to another that is assigned with ’WD repeat’ (i.e. ’WD’ and ’repeat’). In addition, this allows us to aggregate features from the different databases into a homogeneous feature space. For further comparisons, the InterPro entry IDs and signature IDs assigned to the regions are used directly as they require no further processing.
3. **InterPro entry IDs** a collection of unique IDs (which can map to protein families, domain families, repeats, sites), assigned to the regions by InterPro.
4. **Signature IDs** from InterPro member databases a collection of unique IDs (which can map to protein families, domain families, repeats, sites), assigned to the regions by the member databases of InterPro. This also includes unintegrated entries like signal peptide and transmembrane topology predictions from tools like Phobius [29] through InterProScan [17].

The hypothesized advantage of using InterPro entry IDs is that the feature set would be concise and curated, whereas the advantage of using the underlying Signature IDs is that the feature set would be more sensitive to functional differences. This is due to the fact that different databases use different models and methods to classify and identify the assigned features, so not all regions containing the same InterPro IDs will be matched to the same set of signature IDs. For example, the “C2 domain” (IPR000008) groups four contributing signatures and is assigned to both “Fer-1-like protein 5” (A0AVI2) and “Extended synaptotagmin-3” (A0FGR9) proteins. However, only three out of the four signatures matched “Fer-1-like protein 5”, while all four signatures matched “Extended synaptotagmin-3”.

The resulting frequency matrices for the features are then transformed into TF-IDF weights [50] (a well-established technique in Natural Language Processing) to upweight features that occur in fewer proteins and downweight features that occur in many. The dimensions of the different feature types for yeast and human are shown in Suppl. Table S1 and a visual example of the differing pairwise similarity scores (cosine similarity) for a set of 500 regions can be viewed in Suppl. Figure S1.

### 2.3. Our region-specific cost function

To transfer known protein labels to the respective functional regions, we have extended an approach called Group-Instance Cost Function (GICF) [32], initially applied to sentiment analysis of sentences within larger document and user hierarchies. Applied to protein function prediction, these prior works are analogous to identifying positive and negative labels (for a given GO term) for regions within proteins, given only known labels of whole proteins.

This method involves minimizing a cost function that penalizes differences in predicted region scores based on their pairwise feature similarities and also differences between predicted protein scores (aggregated from the constituent region scores) with known protein labels. Additional model components are introduced here to account for differences between predicted region scores with known domain labels (from manually curated databases) and to reduce over-fitting to training data (additional regularization terms have been added).

All together, our cost function consists of 4 terms, each of which enforces the following constraints respectively: (1) regions with similar features should have similar predicted scores, (2) predicted region scores should aggregate to give the correct protein scores when evaluated against known protein labels positive or negative (protein-level constraint),(3) predicted region scores should agree with any known domain-level labels only positive due to the sparsity (region-level constraints), (4) model should not be overfitted to the training set, which is a standard procedure in machine learning but was missing from the original cost function (regularization term).

For each GO term, an independent *θ* is estimated from the following cost function:

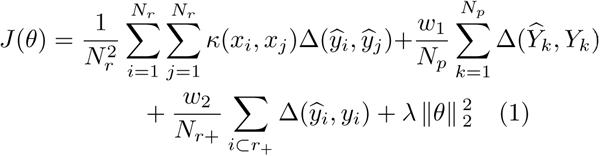

where:

- *y*_*i*_ *∈* {0, 1} is the known label for region *i*,
- *Y*_*k*_ *∈* {0, 1} is the known label for protein *k*,
- 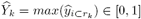 is the predicted score for protein *k*, obtained by getting the maximum predicted score of its containing regions *r*_*k*_,
- *N*_*r*_ is the total number of regions, *N*_*r*+_ is the number of positively annotated regions and *r*_+_ is the subset of positively annotated regions across all proteins,
- *N*_*p*_ is the total number of proteins,
- *κ*(*x*_*i*_, *x*_*j*_) *∈* [0, 1] is the similarity score between regions *x*_*i*_ and *x*_*j*_, calculated using the cosine similarity of the feature vectors,
- Δ is the square loss function, which is the square of the difference between the variables, and
- *w*_*n*_ and *λ* are the trained weights to balance the contributions of the different terms.

The cost function is based on the output of the following logistic regression model:

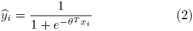

where:

- *x*_*i*_ is the input feature vector for region *i*,
- *θ* is the weight vector of the different contributing features, and
- 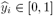 is the predicted score for region *i*.

The choice of logistic regression is due to the binary nature of individual labels (a given protein has the function or it does not) and the resulting probabilistic output that is useful for ranking the predictions. The seed (initial) *θ* was generated for each GO term by fitting a logistic regression model using protein-level features and known protein labels of the same training set. We used minibatch stochastic gradient descent with momentum to train the model. Values of all the hyperparameters are det)ailed in Suppl. Section 3.

### 2.4. Modes of evaluation

### 2.4.1. Region-level evaluation

Due to their sparsity of domain-level annotations and the lack of region-level annotations, there is a lack of gold standard datasets that we can use as a benchmark to effectively evaluate many of the GO terms at the region-specific level. However, to show that this approach can successfully localize GO term labels to the correct regions, we decided to use pairs of binding site annotations from the following databases to evaluate their respective ligand binding GO terms: NBench [38], as a source of nucleic acid binding sites for “DNA binding” (GO:0003677) and “RNA binding” (GO:0003723), and BioLiP [60], as a source of magnesium and zinc ion binding sites for “magnesium ion binding” (GO:0000287) and “zinc ion binding” (GO:0008270). Both databases are semimanually curated from protein structures extracted from the Protein Data Bank and evaluating them in pairs provide convenient sets of negative examples. However, as most proteins do not have structural data for the entire protein length, only regions with at least 80% structural coverage were considered in the evaluation. These selected regions (and their parent proteins) were removed from the training set and used only in the validation and test sets.

For model training, protein-level annotations were obtained from UniProt-GOA and domain-level annotations were obtained from external2GO (see Table 1). Known domain-level annotations were propagated up to the parent proteins for consistency. For validation and testing, a combination of region-level annotations from external2GO and regions containing binding sites were used. Regions from all validation/test proteins are pooled together and evaluated as one set for each GO term using *F*_*max*_, which is the maximum value of the F1 score-harmonic mean of precision and recall. Precision and recall was deemed more appropriate than other evaluation metrics due to the highly skewed class distributions and the F1 score gives us a single value with a convenient threshold.

Due to the very small sample size, the robustness of the rankings was assessed by 100 rounds of random sampling using 50% of the targets as validation (for model selection) and the remaining 50% as test sets. The method is then compared with a whole-protein baseline method (see section 2.4.3) using two-tailed Wilcoxon signed rank test. The details of the region-level datasets are found in Suppl. Section 2.1.

### 2.4.2. Protein-level evaluation

We expanded the number of MF-GO terms tested by conducting a separate protein-level evaluation using a temporal holdout method. Non-IEA (Inferred from Electronic Annotation) annotations at two time-points were used to train and validate/test the model respectively. This ensures a more ’realistic’ approach compared to cross-validation as only proteins with older annotations are used for training, while proteins that gained new annotations are used for validation and testing, allowing for a fully blind test. This also allows us to test if localizing GO term labels to their respective regions can indeed improve overall function prediction at the protein level and if so, for which GO terms. The predicted protein scores are obtained by getting the maximum predicted score of its containing regions. The details of the protein-level temporal holdout datasets are found in Suppl. Section 2.2.

### 2.4.3. Whole-protein baseline method

Logistic regression (implemented using source code from [46]) was used as the **baseline method** for comparison against our method. It is trained directly on features from *whole proteins* (i.e. features from all regions plus those that span multiple regions) with no regularization (*λ* = 0) and the same set of protein-level annotations, with no knowledge of region boundaries at all. The estimated *θ*_*base*_ was also used as the seed input *θ* to our cost function. This would give us an estimate of how well the predictions would do without the constraints of the region-specific framework. As the feature vectors are the same dimensions for regions and proteins, the *θ*_*base*_ trained on whole proteins can be used to predict for both regions and proteins directly, allowing us to compare the scores predicted at the region- and protein-level.

## 3. Results and Discussion

Here, we show the ability of our method to localize binding labels to specific regions within proteins by comparing the predictions directly to binding sites extracted from protein-ligand structures. We also show that the added region-specific framework can lead to improvements in protein-level function predictions for a majority of Molecular Function GO terms that we tested in the section after.

### 3.1. Region-specific localization of binding terms

Here we detail tests of our method over a subset of GO terms can be tied directly to protein sequence via structure, focusing on cases where protein structure analysis provides many unambiguous localizations to many proteins with residue-level resolution.

Results from the region-level evaluation for yeast and human are shown in Figure 2. In each plot, the upper panels show box plots of *F*_*max*_ scores generated over 100 rounds of evaluations for the whole-protein baseline and region-specific methods, while the bottom panels show the differences in median *F*_*max*_ between the two methods for each feature type. All of the differences are significant to .05 level based on test statistics from two-tailed Wilcoxon signed rank test. This head-to-head comparison allows us to eliminate the differences in raw performances based on lack of informative features or annotations, allowing us to focus entirely on the effectiveness of using this region-specific framework for this particular set of feature types and GO terms. The main difference between the models (aside from the region-specific constraints) is that the whole-protein baseline model uses a combination of all the features found within the regions plus additional protein-level features that span multiple regions, like protein families.

**Figure 2:**
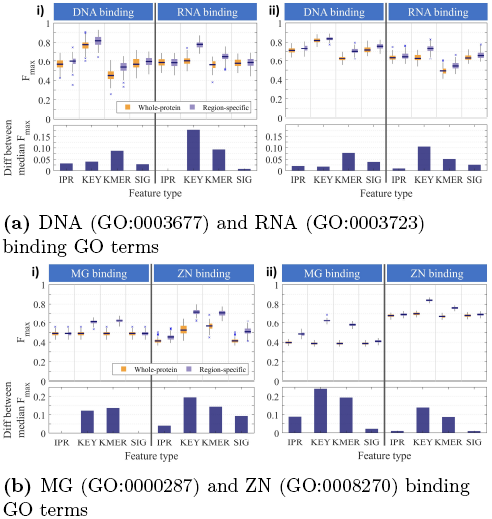
Performance comparisons of region-level predictions using the whole-protein baseline model (orange) versus our region-specific model (purple) for (i) yeast and (ii) human proteomes. KEY = Keywords, KMER = K-mers, IPR = InterPro IDs, SIG = Signature IDs. **Upper panels:** Box plots showing the first quartile (Q1), median, third quartile (Q3) and outliers of maximum *F* measure scores (*Fmax*) generated over 100 rounds of evaluations. **Bottom panels:** Differences between the median *Fmax* scores between methods corresponding to the pair of box plots directly above. Positive values indicate that the region-specific method outperforms the baseline and vice versa for negative values. All differences are significant to .05 level based on test statistics from two-tailed Wilcoxon signed rank test.

From these results, we can see that the performance of our region-specific method is equivalent to or significantly better than the whole-protein baseline across all organisms and feature types for both pairs of binding terms. The biggest gains in region-level performance overall come from **Keywords** and ***K* -mers**, with **Keywords** exhibiting the best raw performance overall. This is not surprising given that both features can be found in regions without domain family assignments and are more discriminating within regions compared to **InterPro IDs** and **Signature IDs**, allowing for greater propagation of labels between regions. More suprisingly, the raw *F*_*max*_ scores for ***K* -mers** are comparable to those of **InterPro IDs** and **Signature IDs** after our region-specific framework is applied, given that *K* -mers are relatively simple representations of the protein sequences and have the smallest dimensions (Suppl. Table S1).

One of the biggest advantages of ***K* -mers** features are that they do not depend on manual, predictive or computationally expensive feature annotation/extraction methods, which means that ***K* mers** feature vectors are guaranteed to be complete and error-free representations of the underlying amino acid sequences. In the future, it would be trivial to extend this method to other, perhaps more informative, sequence-derived representations like ProtVec [15], biophysical properties of the amino acids themselves [13], or even structure-derived representations like contact maps and features extracted from known structures using the Rosetta energy function [2] to take into account shortand long-range interactions between residues.

**Signature IDs** show equivalent or better performance at the region-level compared to **InterPro IDs**, pointing to the value of unintegrated feature annotations, such as signal peptides and transmembrane helices, that would not be included in purely domain-centric methods. InterPro assignments are dominated by Domain and Family entry types (47% and 32% in yeast, 53% and 22% in human, respectively), and in the majority of cases, two regions are either assigned the same InterPro ID, and thus have a similarity score of 1, or are assigned different InterPro IDs, and thus have a similarity score of 0. As shown in Suppl. Figure S1, the resulting pairwise similarity values for **InterPro** are very sparse relative to the other feature types and that sparsity has the effect of reducing the effectiveness of term 1 in the cost function (equation 1), which encourages propagation of labels between regions.

However, the differences in performance between the two feature types are not large and the substantial increase in feature dimension, and thus computational resources required, must be considered. One solution to this problem would be to reduce the dimensions using an autoencoder, an approach that we are also considering as a way to integrate the different feature types into a single, low-dimensional feature space.

### 3.1.1. Structural examples of improved label localization

Here we leverage the very large diversity of RNA and DNA binding proteins with known structures to investigate the performance and resolution of our method. We include cases where sequences lack well annotated domains or where DNA/RNA binding is mediated by additional un-annotated regions (regions not associated with an annotated domain). Figures 3a and 3b show two examples of how DNA and RNA binding GO terms are localized to specific regions within the protein using only **Keywords**. Structures on the left are colored according to predictions made by the whole-protein baseline method, while structures on the right are colored according to predictions made by the region-specific method. Predictions were made at the respective *F*_*max*_ thresholds of each method obtained using the test regions shown in Suppl. Table S2. Shades of green and gray represent positively and negatively labeled regions, respectively, and bound DNA and RNA molecules are shown in black and red, respectively.

**Figure 3:**
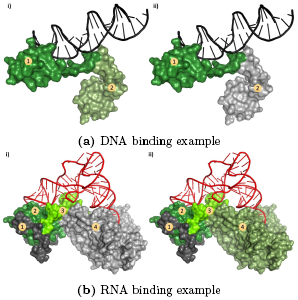
(a) DNA binding predictions for regulatory protein GAL4 (P04386, PDB ID:3coq) and (b) RNA binding predictions for aspartate–tRNA ligase, cytoplasmic (P04802, PDB ID:1asy) from yeast based on (i) baseline and (ii) region-specific methods using **Keywords** features. Structures shown include bound DNA (black) and RNA (red) molecules. **Coloring:** Shades of green (positive) and gray (negative) represent the predicted labels for the given binding GO term at the respective *F*_*max*_ thresholds of each method. **Numbers:** Number labels represent the protein regions ordered from Nto C-terminus. Images are created with PyMOL [52].

Figure 3a shows the crystal structure of part of the “regulatory protein GAL4” (P04386) from yeast. It is composed of two homodimeric chains, one of which is shown here for clarity. Each chain contains a DNA binding and a dimerization region (numbered from Nto C-terminus). Here, we can see that both the DNA binding and dimerization regions are predicted by the whole-protein baseline method to have the DNA binding GO term (both parts colored in green). On the other hand, our region-specific method has correctly removed the DNA binding prediction at the dimerization region below (gray), successfully localizing the GO term only to the direct DNA binding region.

We also looked more closely at the cytoplasmic “Aspartate–tRNA ligase” (P04802), a protein with a high quality RNA-protein experimental structure. Figure 3b shows an RNA binding protein containing four regions (numbered from Nto C-terminus), two of which are associated with known domain families (regions 2 and 4). This protein RNA-complex is from yeast and is also a homodimer with only protein regions forming the protein interface. As before, only one protein chain in this homodimer is shown for clarity. In this case, the whole-protein baseline method (run individually on each region) predicted only regions 2 and 3 to be in contact with RNA, whereas our region-specific method correctly attributed RNA binding to all three regions from 2 to 4. The remaining distal region 1 remains unlabeled, in accordance with lack of protein-RNA contacts observed for this region in the crystal structure.

### 3.2. Expanded functional evaluation at protein level

After the initial proof of concept, which showed that our region-specific method can successfully localize binding GO term labels to specific regions, we extended the analysis to include GO terms from a larger number of functional categories by evaluating the predictions at the protein level. In these results, we show the differences in median *F*_*max*_ performances of our method *relative to* the whole-protein baseline model using the same training, validation and test sets. Figures detailing the individual raw performances for each feature and GO term can be found in the Suppl. section 4.

Figures 4a and 4b show the differences in median *F*_*max*_ of protein-level predictions over all four feature types for yeast and human, respectively. All GO terms are grouped into the shared parental terms (shown in blue on the top left) for clarity.

**Figure 4:**
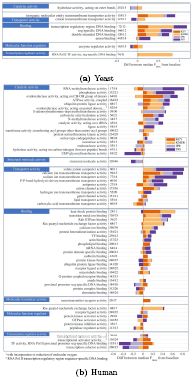
Stacked bar plots showing the differences in performance of protein-level predictions using the best performing region-specific model versus the whole-protein baseline model using non-IEA annotations. KEY = Keywords, KMER = K-mers, IPR = InterPro IDs, SIG = Signature IDs. GO terms are grouped into the parent categories labeled in blue on the top left. The size of each block represents the difference between the median *Fmax* scores of our method and the baseline. Positive (right) means that our method performs better than the baseline and vice versa for negative (left). GO terms are sorted by the absolute sum of the differences across features. Numbers next to each GO term = number of positive training (also includes those inferred from region-level annotations) and testing proteins. All differences shown are significant to at least.05 level.

The differences are shown as colored bars stacked either in the positive (right) or negative (left) direction. Only differences that are significant (p-value < 0.05 level based on two-tailed Wilcoxon signed rank test) are shown on the plot. The numbers next to each GO term represent the number of positive training and testing proteins for that particular GO term (i.e. the particular class distributions during training and testing). As F-scores can be affected by skewed distributions [25], these values are shown here to provide a fuller picture when comparing performances between GO terms, even if the GO terms have been previously filtered to remove very common (>300 occurrences per proteome) and very rare (<10 occurrences per proteome) terms. The number of training examples also include protein annotations inferred from region-level annotations for consistency (since those annotations were used in the cost function during training) and so may be greater than the initial cutoff of 300.

Aside from significant negative correlations (*ρ*=-0.68, p-val=0.04, n=9) between the differences from baseline and the number of positive training and testing proteins for ***K* -mers** in yeast (which is lost when the outlier ’anion binding’ is removed), there were no other notable correlations between the performance differences and the number of positive training or testing proteins for the other feature types in yeast and human.

Our results here suggest that this framework can not only localize GO terms to specific regions, but that it also improves protein-level function predictions for the majority of Molecular Function GO terms evaluated. Furthermore, the performance relative to the whole-protein baseline model can give us a way to characterize region-specificity of different GO terms and thus inform us on how to treat them during the function prediction process.

### 3.2.1. Characterizing region-specificity of GO terms based on feature performance relative to baseline

Overall, the majority of GO terms tested perform significantly better relative to baseline, suggesting that localizing labels to specific regions can improve protein-level predictions as well. In our temporal holdout for yeast, 6 out of 9 MF GO terms showed improvements with our regionspecific model for two or more features. For the human temporal holdout, we see that 41 out of 59 MF GO terms showed improvements, demonstrating that the region-specific framework improves accuracy and coverage of function prediction for more than two thirds of molecular functions. In addition, we can also make conclusions about the ’known’ unit of function specificity (i.e. site, domain family, protein family) for each of the GO terms based on how the different feature types perform relative to baseline.

#### Site-specific GO terms

In both organisms tested, the majority of **Binding** terms show improved performance regardless of feature type, which is not surprising considering the site-specific nature of most binding interactions and the availability of binding site annotations within specific regions. In cases where **InterPro IDs** show significant improvements, it is likely that the predictions are assisted by region-specific InterPro features, such as Repeats and Sites, as they allow labels to be propagated between regions assigned with different domain families. For instance, the GO term “calcium ion transmembrane transporter activity” (GO:0015085) is linked to “Ptype ATPase, phosphorylation site” (IPR018303), while “oxidoreductase activity, acting on CH-OH group of donors” (GO:0016614) is linked to conserved sites like “Short-chain dehydrogenase/reductase, conserved site” (IPR020904) and “Isocitrate/isopropylmalate dehydrogenase, conserved site” (IPR019818).

The importance of non-InterPro entry features, like transmembrane topology and residue annotations, can be seen when **Signature IDs** show much greater improvements than **InterPro IDs**, as is the case for many of the **Transporter activity** terms.

#### Domain-specific GO terms

For GO terms that can be mapped only to InterPro domains, the change in performance of **InterPro IDs** would not be significant but there could be improvements in the other feature types as they may contain other feature annotations that allow for label propagation between regions. For example, performance for GO term “transcription regulatory region DNA binding” (GO:0044212) in yeast improved significantly for all feature types except **InterPro IDs**, which did not show any significant change from baseline. This label is already automatically localized to small regions containing domains like “Zn(2)-C6 fungal-type DNA-binding domain” (InterPro entry: IPR001138), which was not annotated with this GO term at the time of training. The large improvement in ***K* -mers** performance, in particular, suggests that a representation as simple as ***K* -mers** can become very informative when used within our region-specific framework, especially when the GO term can be localized to a small, sequence-specific region.

#### Multidomain GO terms

Significant decreases in performance relative to the baseline across feature types could suggest that the given GO term cannot be localized to specific regions but can be assigned to features spanning multiple regions (e.g. protein families). This could occur if specific features from multiple regions are needed for the association or if the protein family is composed of domains found in diverse proteins, such that a strong association can only be made at the level of the protein family itself.

For example, the GO term “transcription factor activity, RNA polymerase II proximal promoter sequence-specific DNA binding” (GO:0000982) should only be annotated to sequences that contain both the DNA binding and transcriptional activation regions [39]. Individually, DNA binding domains like “Homeobox domain” (IPR001356) and “Zinc finger C2H2-type” (IPR013087) are not sufficient for transcriptional activation and can be involved in a variety of specialized functions: only about 23% and 7% of the training proteins containing these domains are annotated with this specific GO term.

For **InterPro IDs** in particular, if the association to the region features are weak in the training set (i.e. the fraction of proteins with those features that are positively annotated with that GO term is low), then term 1 of the cost function will weaken the associations further due to high region similarity to regions in negatively annotated proteins (refer to a similar discussion in Section 3.1 on sparsity of **InterPro IDs**).

In this case, it is likely that this GO term is too specific to be assigned to individual domains and can only be assigned strictly at the level of the protein family, such as the “E2F family” (IPR015633) and “p53 tumour suppressor family” (IPR002117), where the functional specificity of this GO term is warranted or appropriate. The increases in performance for **Keywords** for these **Transcription regulator activity** terms are due to the enrichment of keywords like ’dna’ and ’homeobox’ in regions of proteins annotated in the test set (7 out of 11 positive proteins), resulting in greater precision and recall even with their weak associations to the GO term itself.

## 4. Conclusion

Here, we have described a pipeline that takes into account regions with unassigned domain families. Our method is built around a cost function that can learn the labels of these regions directly from protein-level annotations and the features of the regions themselves based on the compact biologically-reasonable assumption that functional homology is mediated by regions with similar (if unknown) features.

The results from our region-level evaluation using ligand binding datasets show that our method can successfully localize functions known to be sitespecific to their respective functional regions and performs significantly better than the whole-protein variant across feature types analyzed in both yeast and human. We also evaluate the performance of our region-specific prediction method at the whole-protein level to determine the protein functions that benefit from our explicit region localization and find that, while localization improves performance for some functions, it also decreases performance for others.

This difference in the effect of mapping function to specific regions supports the notion that different GO terms have different levels of operational units and that they should be treated differently in protein function prediction pipelines to take that into account (with some functions tied to small active/binding regions, some tied to domains, and some that need multiple domains and regions for proper functioning). Our results serve as a starting point to begin categorizing GO terms into regionspecific and protein-wide sub groups to maximize the predictive performance of protein function prediction for each GO term and to provide a framework for selecting correct function prediction methods for different functions.

Future work would include introducing a hierarchy of region boundaries within a single protein to allow for different levels of label propagation for different GO terms, and also the use of different feature types simultaneously to consolidate different sources of information. One could, in principle, use autoencoders or NNMF to integrate the different feature types into a single, low-dimensional feature space that would be well suited to our region-level model [21, 35]. We will also experiment with combining our region-specific model with proteinprotein network data to incorporate known overarching relationships between proteins for a more comprehensive function prediction tool.

## Acknowledgements

We thank Vladimir Gligorijević, Meet Barot and Noah Youngs for their assistance and discussions that were instrumental to the work. This work was supported in part through the NYU IT High Performance Computing resources, services, and staff expertise in New York City and Abu Dhabi.

## Supplementary Material

### 1 Feature representations

**Table S1:** Dimensions of features

| Organism | *K*-mers | Keywords | InterPro IDs | Signature IDs |
| --- | --- | --- | --- | --- |
| Yeast | 1,728 | 7,423 | 5,847 | 15,451 |
| Human | 1,728 | 16,968 | 13,394 | 41,596 |

Number of unique features that are present in at least 1 protein in the given organism. For Keywords, it must be present in at least 2 proteins to remove the majority of misspellings.

**Figure S1:**
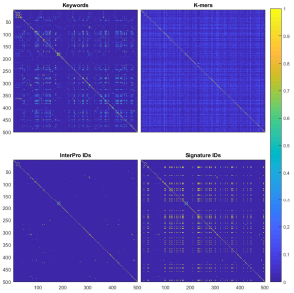
Distribution of pairwise cosine similarity scores between 500 regions for different features.

### 2 Train/validation/test sets

#### 2.1 Region-level evaluation

**Table S2:** Number of train, validation/test regions/proteins used in the region-level validation of our method.

|  |  | Regions | Proteins |
| --- | --- | --- | --- |
| Ligands | Count type | #(train, valid/test) | #(train, valid/test) |
| **Yeast** |  |  |  |
| DNA/RNA | Total<sup>1</sup> | (9712, 183) | (3257, 70) |
|  | DNA binding<sup>2</sup> | (134, 52) | (268, 27) |
|  | RNA binding<sup>2</sup> | (179, 72) | (338, 38) |
| MG/ZN | Total<sup>1</sup> | (9316, 606) | (3131, 266) |
|  | MG binding<sup>2</sup> | (42, 197) | (34, 157) |
|  | ZN binding<sup>2</sup> | (133, 157) | (103, 130) |
| **Human** |  |  |  |
| DNA/RNA | Total<sup>1</sup> | (37878, 301) | (9272, 154) |
|  | DNA binding<sup>2</sup> | (531, 138) | (856, 97) |
|  | RNA binding<sup>2</sup> | (114, 98) | (335, 65) |
| MG/ZN | Total<sup>1</sup> | (35153, 2287) | (8751, 1154) |
|  | MG binding<sup>2</sup> | (31, 545) | (79, 459) |
|  | ZN binding<sup>2</sup> | (451, 1159) | (402, 760) |

<sup>1</sup>Total number of regions/proteins in set.
<sup>2</sup>Number of positive binding regions/proteins.

#### 2.2 Protein-level evaluation

The protein set was divided as follows into training, validation and test sets. See Figure S2 for a visual reference.

- **Training set** - Proteins that had at least 1 annotation at older time-point and did not gain new annotations by new time-point;
- **Validation set** - Proteins that had at least 1 annotation at older time-point and gained new annotations by new time-point;
- **Test set** - Proteins that did not have annotations at older time-point but gained new annotations by new time-point.

After the protein sets have been established, GO terms that fit the following criteria were selected to be tested with the model:

- Have between 10 and 300 positive annotations in the training set;
- Have at least 10 positive annotations in both the validation and test sets.

These constraints were used to ensure that there is sufficient positive annotations to train and test the model with in order to get meaningful performance reports.

**Figure S2:**
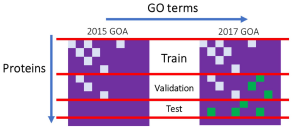
Schematics of temporal holdout data splitting. Matrices: Rows represent proteins, columns represent GO terms. Small boxes: GO terms assigned to proteins. Gray = present in the older (e.g. 2015) Gene Ontology Annotation (GOA) database. Green = present in the newer (e.g. 2017) GOA database.

**Table S3:** Number of train, validation and test proteins used in the protein-level temporal holdout validation of our method.

| Yeast |  | Human |  |
| --- | --- | --- | --- |
| GO terms<sup>1</sup> | #(train, valid, test) | GO terms<sup>1</sup> | #(train, valid, test) |
| 9 | (3302, 1083, 250) | 59 | (9347, 4266, 1862) |

<sup>1</sup> Total number of MF-GO terms evaluated. Parental GO terms that also satisfy selection criteria were removed.

### 3 Model hyperparameters

All hyperparameter values are shown in Table S4. We optimize the cost function (Eqn. 1 in the main manuscript) using minibatch stochastic gradient descent (SGD) `optimizer` with `momentum`. To prevent overfitting, we implemented `early stopping` by monitoring the *F*_*max*_ performance of the training set after every 50 iterations. We stop the training when the variance of the last 50 *F*_*max*_ values drops below the minimum variance threshold and after the minimum number of iterations has been completed.

The model is trained with a batch size of at least 128, depending on the number of positive examples for the particular GO term. This is due to the sparsity of positive annotations and the need to have at least one positive example in each minibatch to calculate the *F*_*max*_ for the stopping criteria. We generated the minibatches using a stratified sampling strategy to ensure that the class distribution is maintained during training as it appears to perform slightly better overall compared to random sampling in preliminary tests.

We also decreased the learning rate based on the epoch cycle using the given exponential decay formula to improve rate of convergence.

The range of *ω* _1_, *ω* _2_ and *λ* values used in the cost function are shown under ’Cost function’ and they were narrowed down using a grid search on the region-level evaluation dataset.

**Table S4:** Model hyperparameters.

| Hyperparameters | Values |
| --- | --- |
| Stochastic gradient descent |  |
| batch_size | $\max \left( 128, \text{ceiling} \left( \frac{\text{number of training examples}}{\text{number of positive examples}} \right) \right)$ |
| max_epochs | 500 |
| min_iteration | 500 |
| lr | 0.5 |
| lr_decay | $(1 - 0.1)^{\text{floor}(\frac{\text{epoch}}{2})}$ |
| momentum | 0.9 |
| min_variance | 0.005 |
| Cost function |  |
| $w_1$ range | [1e-3 1e-2 1e-1 1e0 1e1] |
| $w_2$ range | [1e-3 1e-2 1e-1 1e0 1e1] |
| $\lambda$ range | [1e-5 1e-4 1e-3 1e-2 1e-1] |

### 4 Temporal holdout performance

#### 4.1 Yeast proteome

**Figure S3:**
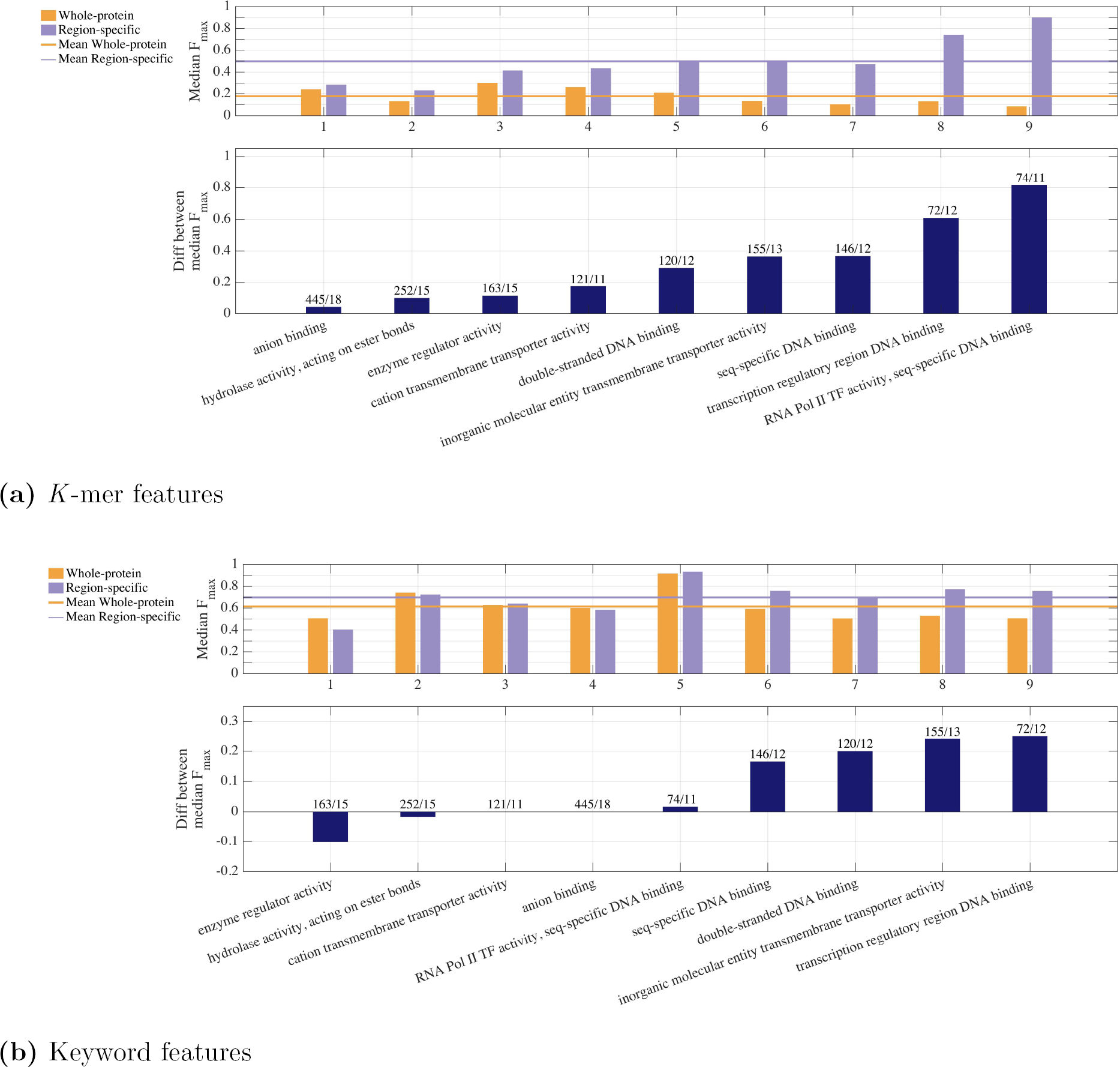
Performances of protein-level predictions on 9 Molecular Function GO terms using *K* -mer and Keyword feature types for yeast proteome on test set using non-IEA annotations only. Median *F*_*max*_ scores for the baseline model and our best-performing model for each GO term tested are shown as bar plots in the upper panel. The GO terms are sorted in ascending order of their differences, which are shown in the panel directly below. Positive differences mean that our method performs better than the baseline and negative differences mean the opposite.

**Figure S3:**
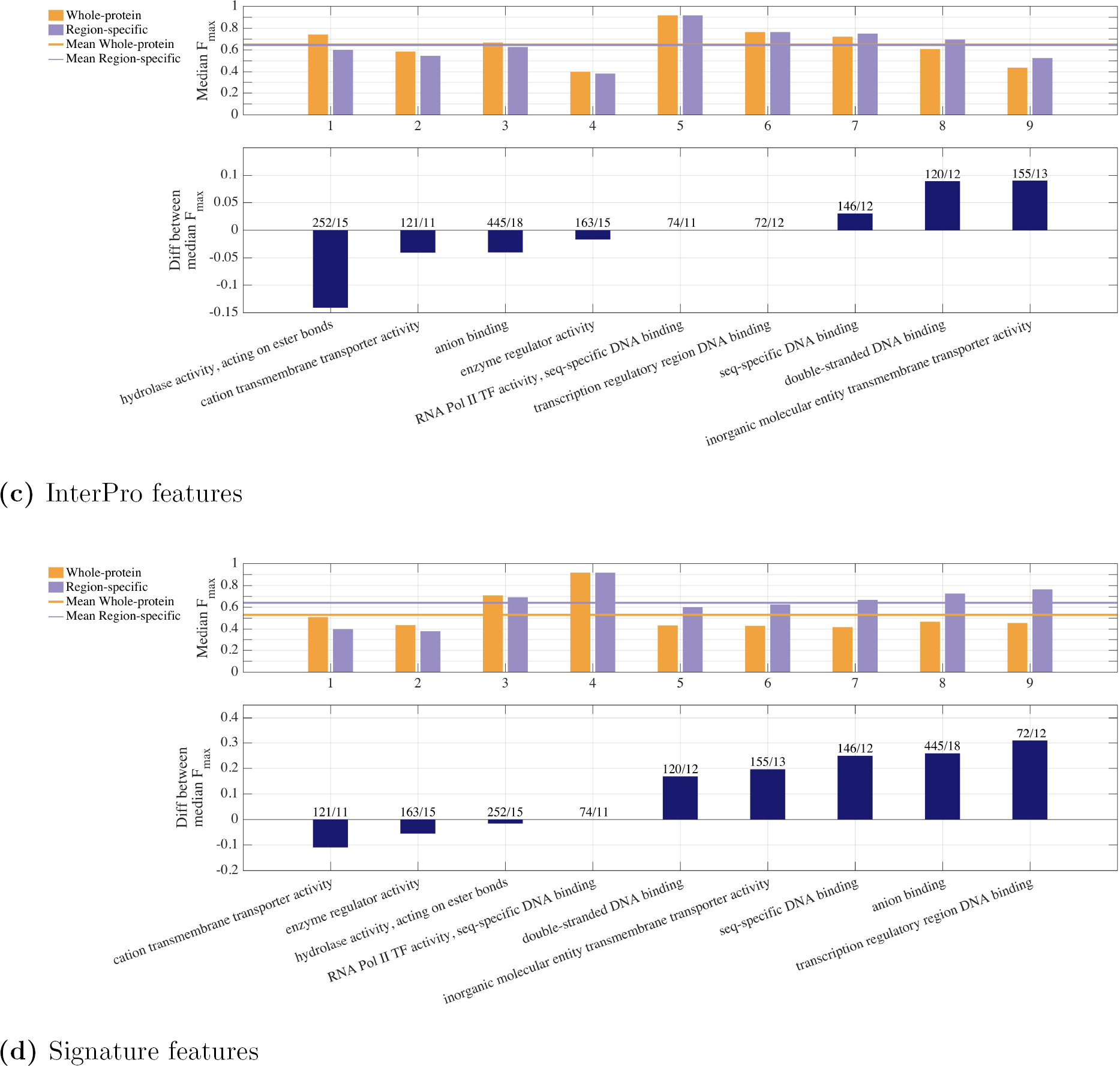
Performances of protein-level predictions on 9 Molecular Function GO terms using InterPro and Signature feature types for yeast proteome on test set using non-IEA annotations only. Median *F*_*max*_ scores for the baseline model and our best-performing model for each GO term tested are shown as bar plots in the upper panel. The GO terms are sorted in ascending order of their differences, which are shown in the panel directly below. Positive differences mean that our method performs better than the baseline and negative differences mean the opposite.

#### 4.2 Human proteome

**Figure S4:**
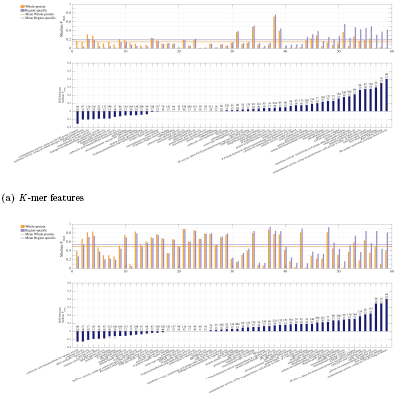
Performances of protein-level predictions on 59 Molecular Function GO terms using *K* -mer and Keyword feature types for human proteome on test set using non-IEA annotations only. …* = RNA Pol II transcription regulatory region seq-specific DNA binding. Median *F*_*max*_ scores for the baseline model and our best-performing model for each GO term tested are shown as bar plots in the upper panel. The GO terms are sorted in ascending order of their differences, which are shown in the panel directly below. Positive differences mean that our method performs better than the baseline and negative differences mean the opposite.

**Figure S4:**
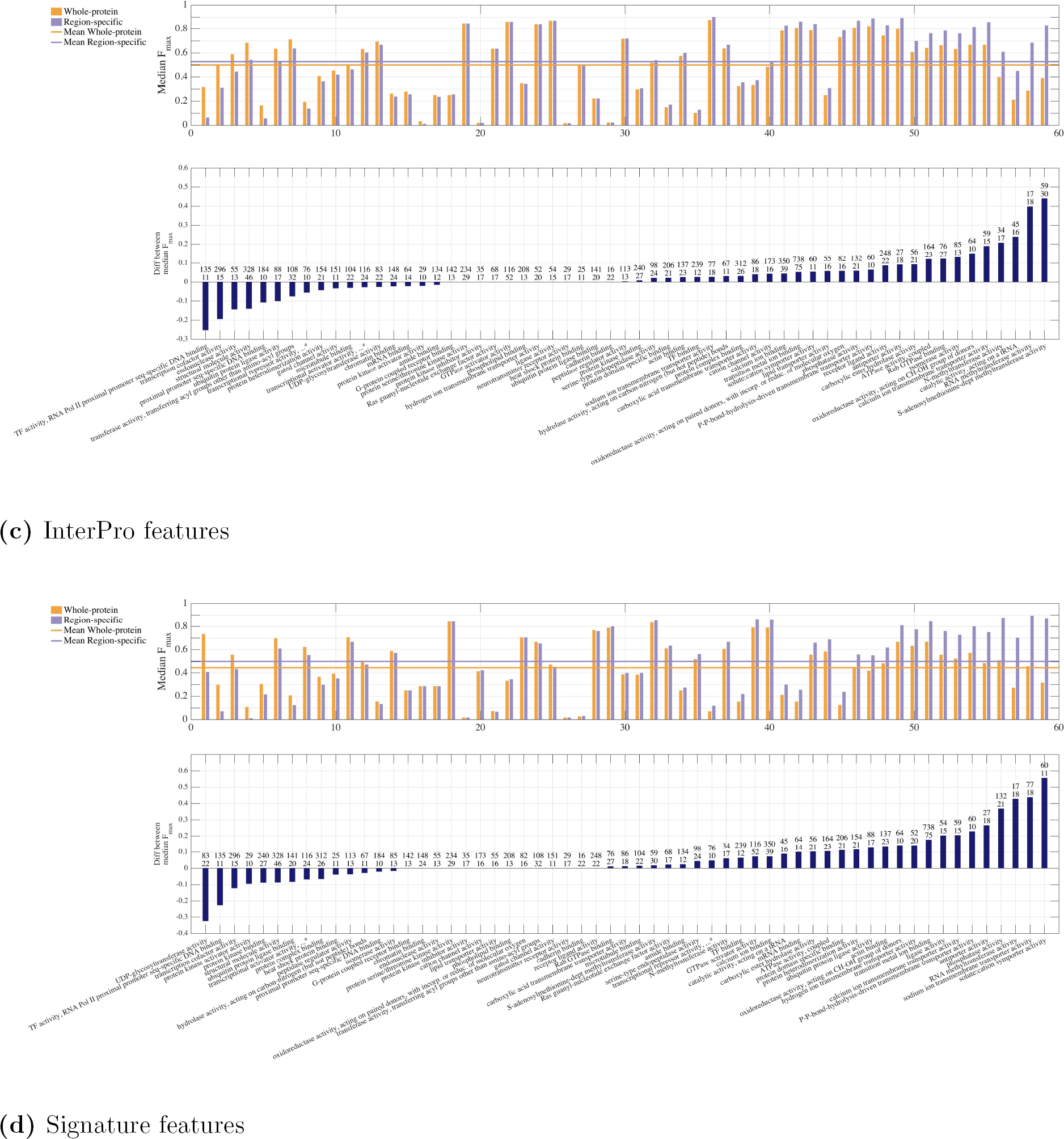
Performances of protein-level predictions on 59 Molecular Function GO terms using InterPro and Signature feature types for human proteome on test set using non-IEA annotations only. …* = RNA Pol II transcription regulatory region seq-specific DNA binding. Median *F*_*max*_ scores for the baseline model and our best-performing model for each GO term tested are shown as bar plots in the upper panel. The GO terms are sorted in ascending order of their differences, which are shown in the panel directly below. Positive differences mean that our method performs better than the baseline and negative differences mean the opposite.

